# The human ApoE4 variant reduces functional recovery and neuronal sprouting after incomplete spinal cord injury in male mice

**DOI:** 10.1101/2020.11.05.369900

**Authors:** Carlos A. Toro, Jens Hansen, Mustafa M. Siddiq, Kaitlin Johnson, Wei Zhao, Daniella Azulai, Dibash K. Das, William Bauman, Robert Sebra, Dongming Cai, Ravi Iyengar, Christopher P. Cardozo

## Abstract

Spinal cord injury (SCI) is a devastating form of neurotrauma. Patients who carry one or two ApoE4 alleles show worse functional outcomes and longer hospital stays after SCI but the cellular and molecular underpinnings for this genetic link remain poorly understood. Thus, there is a great need to generate animal models to accurately replicate the genetic determinants of outcomes after SCI to spur development of treatments that improve physical function. Here, we examined outcomes after a moderate contusion SCI of transgenic mice expressing human ApoE3 or ApoE4. ApoE4 mice have worse locomotor function and coordination after SCI. Histological examination revealed greater glial staining in ApoE4 mice after SCI associated with reduced levels of neuronal sprouting markers. Bulk RNA sequencing revealed that subcellular processes (SCPs), such as extracellular matrix organization and inflammatory responses, were highly-ranked among upregulated genes at 7 days after SCI in ApoE4 variants. Conversely, SCPs related to neuronal action potential and neuron projection development were increased in ApoE3 mice at 21 days. In summary, our results reveal a clinically relevant SCI mouse model that recapitulates the influence of ApoE genotypes on post-SCI function in individuals who carry these alleles and suggest that the mechanisms underlying worse recovery for ApoE4 animals involve glial activation and loss of sprouting and synaptic activity.

## INTRODUCTION

Spinal cord Injury (SCI), one of the most devastating forms of neurotrauma, results in life-long disabilities and loss of independence. SCI affects approximately 300,000 individuals in the United States with more than 17,000 new cases each year (National Spinal Cord Injury Statistical, 2020). SCI interrupts communication between the brain and the tissues innervated by motor and sensory neurons that emerge below the injury resulting in paralysis, loss of sensation and dysregulation of the autonomic nervous system (Lang et al., 2015). Despite successful efforts toward reducing complications following SCI (O’Shea et al., 2017), there are no cell-based or pharmacologic approaches that consistently improve sensory or locomotor function after SCI in humans. Additional knowledge with regard to the cellular and molecular mechanisms that determine functional recovery is essential to stimulate further research to develop and test interventions that will improve function after SCI. Moreover, the understanding of genetic factors that may influence outcomes after SCI, such as the ability to walk, grasp or lift, are profoundly under investigated but may provide new insights with regard to the cellular and molecular determinants of outcomes after SCI that determine functional recovery. One important genetic factor to consider is Apolipoprotein E (ApoE), a well-characterized lipoprotein involved in cholesterol trafficking and lipid metabolism (Huang and Mahley, 2014;Arenas et al., 2017). In the central nervous system (CNS), ApoE is mainly synthesized in and secreted by astrocytes, and evidence from human studies and animal models is consistent with the notion that ApoE is a key determinant of the response to different types of CNS injury (Poirier, 1994;Teasdale et al., 1997).

In humans, there are three common natural variations of the gene that encodes ApoE. These are known as: ApoE2, ApoE3 and ApoE4 and have a worldwide frequency of 8.4-, 77.9-, and 13.7% respectively (Liu et al., 2013). These variants encode for proteins that differ at just one or two amino acid residues (Mahley and Rall, 2000) yet have very dissimilar physiological actions and are linked to the clinical outcomes of several medical conditions (Saunders et al., 1993;Teasdale et al., 1997;Laskowitz et al., 1998;Friedman et al., 1999;Jha et al., 2008;Sun et al., 2011). Mice on the other hand, express only a single ApoE variant; its replacement with any of the human isoforms will also generate different physiological outcomes in response to injury or stress. Consequently, these targeted replacement (TR) mouse models that carry human ApoE variants are increasingly being used to study how ApoE alleles contribute to health and disease within the CNS (Fryer et al., 2005;Castellano et al., 2011).

ApoE plays an active role in axonal growth, synaptic formation and remodeling (Xu et al., 2014) and the clearance of apoptotic bodies during neuronal apoptosis (Elliott et al., 2007). Several studies have reported a significant upregulation of ApoE following severe neurological damage such as SCI (Seitz et al., 2003) and traumatic brain injury (TBI) (Iwata et al., 2005). Interestingly, the presence of the ApoE4 isoform is directly associated with worse outcomes after mild and severe TBI (Smith et al., 2006;McKee and Daneshvar, 2015), is related with early mortality (Jordan et al., 1997), extended coma (Friedman et al., 1999), and is the only known genetic risk factor for sporadic Alzheimer’s Disease (Rebeck et al., 1993). Moreover, clinical studies have examined the possibility that ApoE variants impact functional outcomes after SCI and revealed that the presence of the ApoE4 allele is strongly correlated with significantly worse neurological outcomes and longer hospital stays (Jha et al., 2008;Sun et al., 2011). ApoE4 carriers show less motor recovery during rehabilitation than individuals with other ApoE alleles, as determined by applying the American Spinal Injury Association (ASIA) motor score during rehabilitation (Jha et al., 2008). Despite clinical evidence showing ApoE variants as genetic factors that modulate outcomes after SCI, the cellular and molecular mechanisms by which they influence functional recovery are not understood.

The spontaneous partial recovery of function observed after SCI occurs through neuroplasticity whereby surviving neurons extend projections, forming new, functional neural circuits that bridge the injury. Neuroplasticity is thought to arise through sprouting of axon branches from descending fibers and projection of dendrites and axons from propriospinal neurons in an attempt to connect descending fibers with lower motor neurons; it is also likely that there are cellular changes that strengthen the synaptic connections that remain after trauma or form as a result of spouting (Hutson and Di Giovanni, 2019). Sprouting is followed by pruning, and both events have been documented and provide a logical basis for understanding the cellular processes obligate for neuroplasticity (Goldstein et al., 1997). Other studies have provided compelling evidence of the formation of functioning propriospinal neural circuits that bridge the lesion site after SCI and presumably arise via sprouting (Courtine et al., 2008). Despite the lack of molecular insights as to how ApoE variants influence outcomes after SCI, several studies have suggested that ApoE4 reduces the effectiveness of sprouting. *In vitro* research using neurons from mice expressing human ApoE4 showed impaired neurite outgrowth as compared to those carriers of ApoE2 or ApoE3 variants (White et al., 2001;Seitz et al., 2003). Moreover, ApoE4 mice have been reported to show reduced sprouting after an entorhinal lesion (White et al., 2001).

The influence of ApoE genotype on neuroplasticity, neuronal regeneration, secondary injury or glial reaction after SCI are unknown. To identify the effects of ApoE variants on outcomes after SCI, we administered a moderate severity mid-thoracic contusion SCI to TR ApoE3 or ApoE4 mice and examined locomotor function as well as histologic markers of neuroplasticity and glial reaction, and changes in gene expression patterns as assessed by RT-qPCR and bulk RNA-seq. Our findings revealed that, after SCI, TR ApoE4 mice show less functional recovery, which is associated with less sprouting and greater glial reaction. Moreover, transcriptomic profiling of the lesion site suggests that the initial response in ApoE4 mice is characterized by inflammation and collagen turnover when compared to ApoE3 mice. Transcriptomic profiles obtained at day 21 post SCI suggest that the reduced potential for neuroplasticity and neuroregeneration observed in ApoE4 mice is manifested at the mRNA level as differences in enriched subcellular processes related to neuronal action potential, neurotrophin signaling and neuron projection development for TR ApoE3 in comparison to TR ApoE4 SCI mice.

## MATERIALS AND METHODS

### Animals

All use of live animals was approved by the Institutional Animal Care and Use Committee at James J. Peters Veterans Affairs Medical Center (JJP VAMC) and was conducted in accordance with PHS Policy on Humane Care and Use of Laboratory Animals and the Guide. Human ApoE3 and ApoE4 knock-in mouse models (RRID: IMSR_TAC:1549 and 2542), in which the murine ApoE gene was replaced with the coding region for either human ApoE variant (Grootendorst et al., 2005;Wang et al., 2005;Rodriguez et al., 2013), were purchased from Taconic Biosciences Inc., bred at JJP VAMC and housed in a room with controlled photoperiod (12/12 h light/dark cycle) and temperature (23–25°C) with ad libitum access to water and pelleted chow.

### Experimental Design

All mice were genotyped by qPCR before weaning using a small tissue sample from the tip of the tail. At 3-to 5-months of age, animals weighing between 25.3 and 25.9 grams for males (Supplementary Figure 1A) and 23.7 and 24.5 grams for females (Supplementary Figure 1B) were randomly assigned to SCI or sham-operated groups (Table 1). Experimental design and timeline are provided (Figure 1).

**Table 1.** Groups numbers/procedures Experimental Group Numbers

| Table 1. Experimental Group Numbers |  |  |  |
| --- | --- | --- | --- |
| Genotype/Procedure | Timepoint |  |  |
|  | 7-dpi | 14-dpi | 28-dpi |
| ApoE3 / Sham | N= 10M / 10 F | N= 10 M / 10 F | N= 10 M / 10 F |
| ApoE4 / Sham | N= 10 M / 10 F | N= 10 M / 10 F | N= 10 M / 10 F |
| ApoE3 / C-SCI | N= 12 M / 12 F | N= 12 M / 12 F | N= 14 M / 16 F |
| ApoE4 / C-SCI | N= 12 M / 12 F | N= 12 M / 12 F | N= 14 M / 16 F |

**Figure 1.**
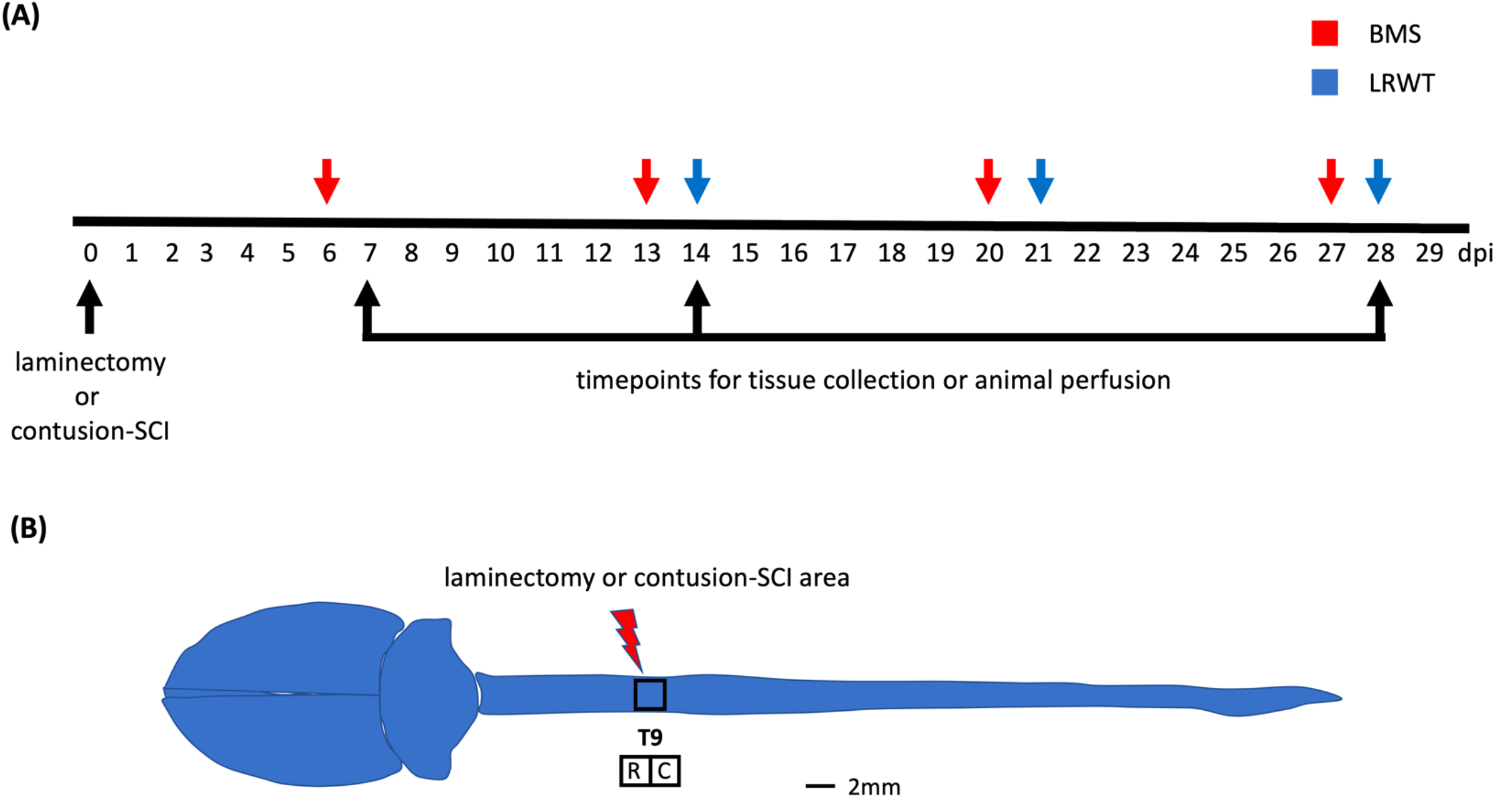
Behavioral examination and anatomical site used for surgical procedures. (A) Timeline of surgeries, behavioral testing and timepoints for tissue collection and animal perfusion. BMS: Basso mouse scale; LRWT: ladder rung walk test. (B) Mouse brain and spinal cord, and site of laminectomy or contusion-SCI at thoracic vertebrae 9 (T9). Adapted from Bacskai et. al, 2013 (Bacskai et al., 2013). Upper black square represents the area for laminectomy-only or contusion-SCI. Lower squares represent segments collected for biochemical studies and for RNA-seq. R: rostral; C: caudal. Scale bar, 2 mm.

### Spinal Cord Injuries

An incomplete SCI was performed after induction of anesthesia by inhalation of 3% isoflurane. Briefly, anesthetized animals were placed on warming blankets heated by recirculating water at 37° C and hair was removed over the thoracic spine. After cleaning the area, sham and SCI animals underwent a laminectomy to expose the dura at thoracic vertebrae 9 (T9). The size of the opening of the vertebral column created by the laminectomy was approximately 2 mm in diameter, was consistent among animals and gave enough room to allow the impact probe to reach the dura without touching bone, muscles or other surrounding tissues and thus produce a controlled, reproducible injury. The lateral part of the vertebrae surrounding area of the laminectomy site containing the facets was carefully preserved in order to maintain vertebral column stability. Muscles on the lateral processes of vertebrae T8 to T10 were peeled away from bone by blunt dissection permitting the stabilization of the vertebral column using forceps attached to the Infinite Horizons (IH) clamping platform. For the SCI groups (Table 1), the IH impactor was used to strike and displace the exposed cord through a computer driven probe connected to a force transducer (Scheff et al., 2003). Both, ApoE3 and ApoE4 animals received a 50 kdyne contusion SCI. The impacting tip was centered from right to left above the exposed cord over the center of the T9 vertebral body to generate an injury affecting the left and right hindlimbs equally. The lesion was confirmed by noting, under a magnifier, symmetric bruises on both sides of the dorsal median sulcus. Parameters obtained from the IH device, such as impact force and cord displacement were recorded for each animal (Supplementary Figure 1C,D). Muscle was later sutured, and skin was closed using 7-mm wound clips. After surgery, mice were placed in cages with Alpha-Dri bedding (Newco Distributors, INC.) and allowed to recover on heating pads warmed with recirculating water for 72 hours. All animals received pre-warmed lactated Ringers solution (LRS), carprofen and Baytril post-operatively until fully recovered. Wound clips were removed after 10 days. Urine was manually expressed twice daily by pressing on the bladder with gentle massage until spontaneous voiding was observed; animals were checked for retained urine at least daily for the length of the study.

### Group Numbers and Exclusion Criteria

A summary of the number of mice for each experimental group is shown (Table 1). The criteria for excluding animals from the study included: i) detection, under a magnifier, of asymmetric bruising of the cord during laminectomy, ii) awareness of manual slippage at the clamping platform when stabilizing the vertebral column with the IH device forceps, iii) bone impact, as noted by irregular time versus force curves on the IH software platform and a much higher than expected actual impact force, and iv) poor or better than expected performance on behavioral tests at 6-days post injury (dpi), reflected by data values three standard deviations below or above the group mean. In addition, other criteria for excluding spinal cord tissues after euthanasia included: i) faulty animal/tissue perfusion, and ii) failure to maintain the integrity of the tissue for vibratome processing.

### Behavioral Testing

Functional recovery after laminectomy/SCI was tested using the Basso Mouse Scale (BMS) open-field test (Basso et al., 2006), and the ladder rung walk test (LRWT) (Cummings et al., 2007), at the time points specified (Figure 1A). The BMS locomotor score is a well-established method for evaluating the severity of impairments in locomotor function after SCI, whereas the LRWT is useful for evaluating coordinated stepping and fine motor skills. The BMS open-field locomotor function test was performed in a commercial 36-inch diameter kiddie pool with a plexiglass bottom. Animals were recorded with a GoPro camera while being observed moving about the pool. BMS locomotor function was scored on a 9-point scale by blinded observers, as described previously (Basso et al., 2006). For LRWT scores, animals were placed at one end of a commercially available ladder for mice and observed as they attempt to explore and cross the ladder (Cummings et al., 2007). Briefly, the apparatus consists of a horizontal ladder beam with 74 rungs suspended 20 inches above the surface with escape boxes at both ends of the ladder. The ladder has 4 mm diameter rungs spaced 12 mm apart. The animals were recorded using a GoPro camera located under the ladder that was moved manually to keep the animal in-frame. Videos were reviewed by two blinded observers who recorded the number of correct steps and errors. Animals were familiarized with the kiddie pool and LWRT apparatus a week prior the surgeries. Animals were also gently handled and scruffed during the week before surgeries to familiarize them with staff and the sensations of being handled.

### Tissue Harvest

At time of euthanasia (7-, 14-, or 28-dpi), 3 to 5 animals per group (Table1) were randomly selected to undergo perfusion-fixation under a deep surgical plane of anesthesia that was achieved using 100 mg/kg of ketamine and 30 mg/kg of xylazine intraperitoneally. Mice were euthanized by transcardial perfusion with sterile saline then fixed using ice-cold 4% paraformaldehyde (PFA) in 0.1 M phosphate-buffered saline (PBS). After perfusion, spinal cords were removed and post-fixed in 4% PFA for 48 h, transferred to sterile PBS and stored at 4 C until sectioning. Fresh spinal cord tissues from the remaining animals of each group (Table 1) were collected after inducing deep anesthesia achieved by inhalation of 3% isofluorane followed by decapitation. Either a block 4 mm in length containing the injury epicenter or ∼2 mm sections rostral and caudal from the epicenter, were collected and snap frozen in liquid nitrogen then stored at −80 C for biochemical analysis; tissues from comparable areas of spinal cords of sham-controls were also collected. A diagram of mouse spinal cord and segments dissected are depicted in Figure 1B.

### Immunofluorescence Staining

Transverse 30-micron sections of perfusion-fixed male spinal cords were cut with a vibratome (Leica) and used for immunofluorescence staining for proteins of interest. Details regarding antibodies used for these studies are shown in Table 2. Immunostaining was performed for: i) GAP-43 (dil 1:500, Abcam, #ab16053), a marker of regenerating axons used to identify sprouting; ii) growth cone formation by using 2G13 (dil 1:500, Novus # NB600-785); iii) presynaptic vesicles using synaptophysin (dil 1:500, Novus # NBP2-25170); iv) astrocyte activation through staining with GFAP (dil 1:500, Abcam #ab7260); and macrophages and microglia by labeling IBA-1(dil 1:500, Abcam # ab178846), as previously described (Perez-Garcia et al., 2018). Secondary antibodies Alexa Fluor 488-conjugated goat anti-mouse IgG (Abcam # ab150113); and Alexa Fluor 647-conjugated goat anti-rabbit IgG (Abcam # ab150079) were used at a dilution of 1:2500. Immuno-stained sections were imaged with a Zeiss 700 confocal microscope (Carl Zeiss). For quantitative measurements, representative injured spinal cord images from four SCI male mice per genotype were captured. Blinded quantification was performed using ImageJ software (version 2.1.0/1.53c, National Institute of Health, USA). Integrated density of pixels was measured in representative areas of the ventral horn from each section, as previously described (Oliveira et al., 2004). The integrated pixel density was calculated for each section of spinal cord, and then a mean value for each spinal cord was calculated. The data are represented as the mean ± SEM. The fluoromyelin staining was performed according to the manufacturer’s protocol (FluoroMyelin green, ThermoFisher).

**Table 2.**
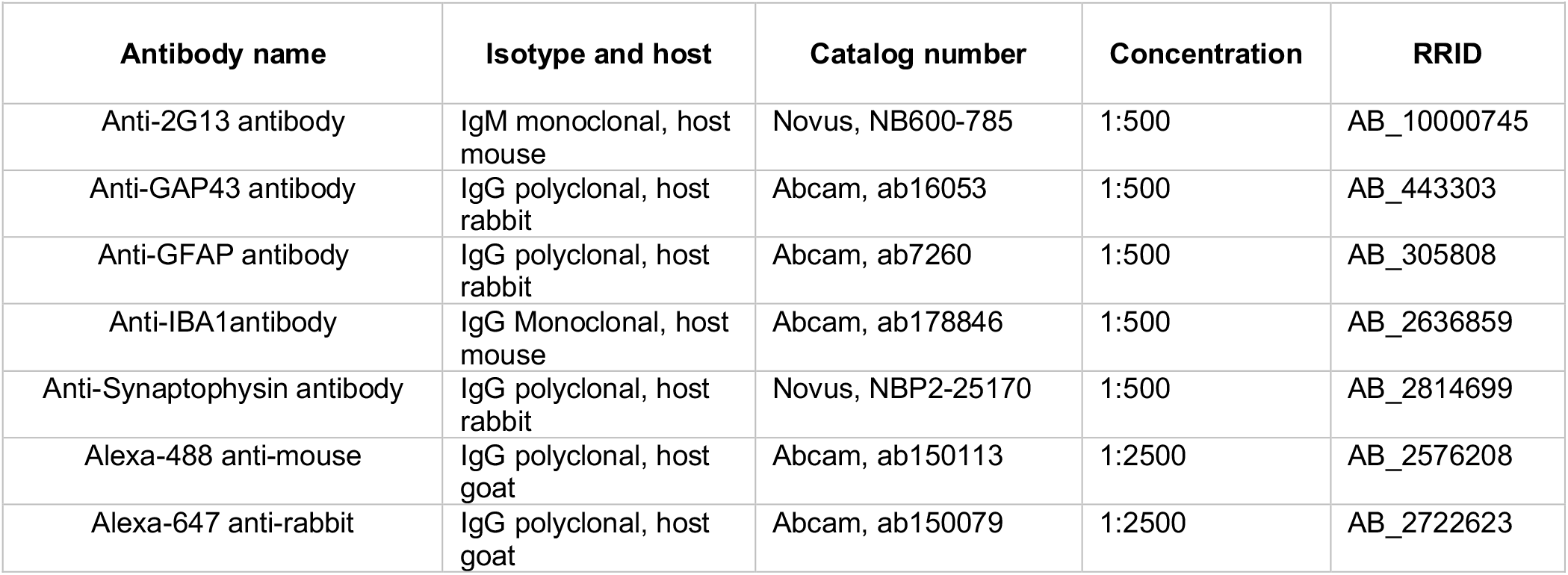
Antibodies Antibodies for immunohistochemistry

### RNA Extraction, Reverse Transcription and Qpcr

Total RNA was extracted from spinal cord segments. TRIzol reagent (ThermoFisher) was used to extract total RNA from samples from shams and SCI ApoE3 and ApoE4 male mice collected at 14-dpi following the manufacturer’s instructions and methods previously described (Toro et al., 2018). Total RNA concentrations were determined by absorbance at 260 nm using a Nanodrop spectrophotometer (Thermo Scientific), and later transcribed into cDNA in a volume of 20 μl using Omniscript reverse transcriptase (Qiagen). The mRNAs of interest were measured using the PowerUp SYBR Green Master Mix (Thermofisher). Primers (Table 3) were designed with help of the Primer Blast program from NCBI. All PCR reactions were carried out using a QuantStudio 12K Real-Time PCR system (Life technologies) in a 10 μl volume, with 1 μl of cDNA, 1 μl of primers (10 μM stock), 3 μl of molecular grade H20, and 5 μl of SYBR Green. The PCR reaction was performed using the following conditions: 95 °C for 5 min, followed by 40 cycles of 15 s at 95 ° C and 60 s at 60 °C. Formation of a single SYBR Green-labeled PCR amplicon was verified by running a melting curve analysis. Threshold cycles (CTs) for each PCR reaction were identified by using the QuantStudio 12K Flex software. To construct standard curves, serial dilutions were used from 1/2 to 1/512 of a pool of cDNAs generated by mixing equal amounts of cDNA from each sample. The CTs from each sample were compared to the relative standard curve to estimate the mRNA content per sample; the values obtained were normalized for procedural variations using peptidylprolyl isomerase A (Ppia) mRNA as the normalizing unit.

**Table 3.**
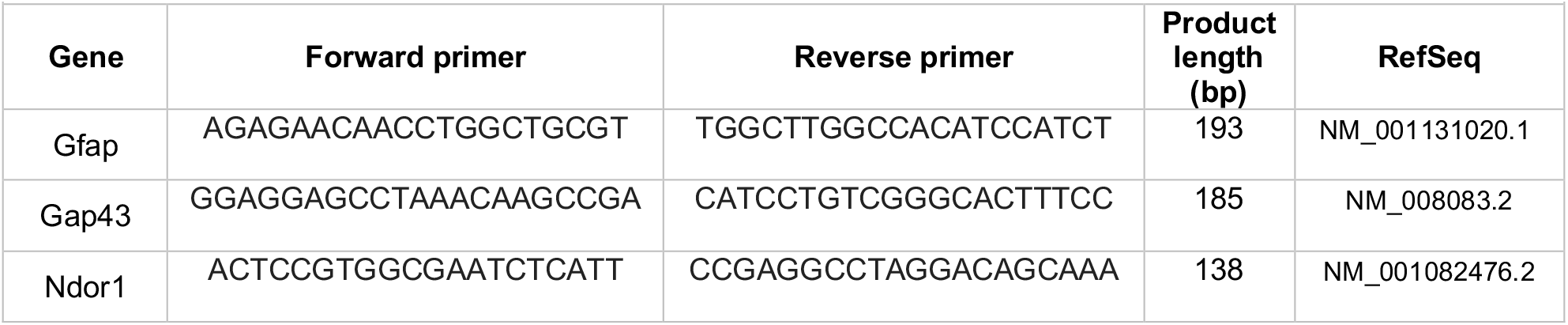
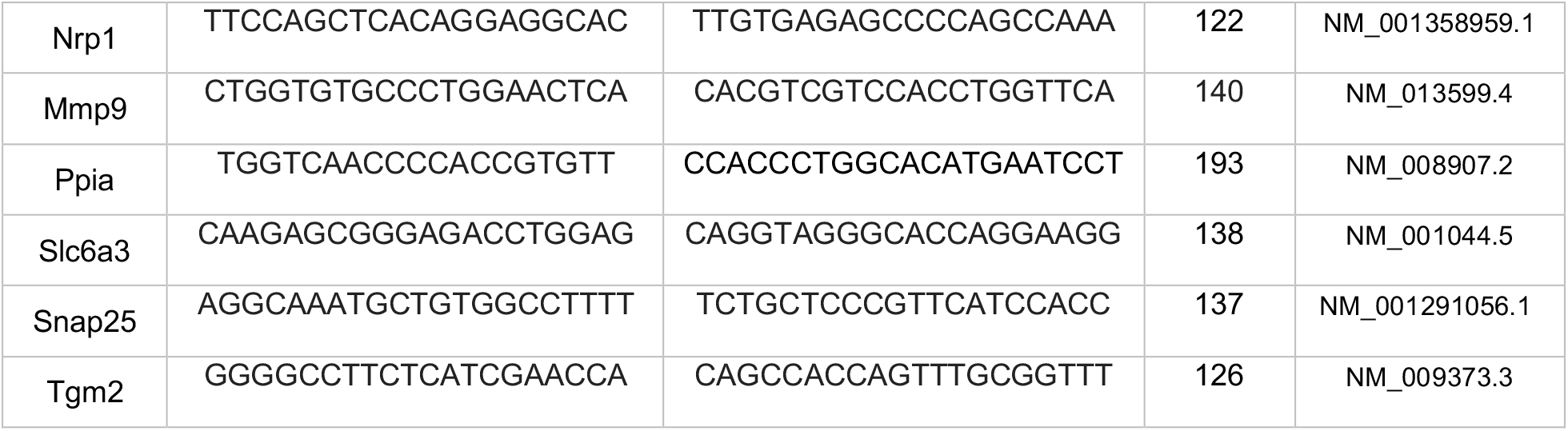
qPCR Primers Primer sequence for RT-qPCR

### Transcriptomic profiling by RNA sequencing

Total RNA was extracted, as described above, from spinal cord segments containing the lesion site (∼4 mm), from male mice at 7-and 21 days post SCI (N=3 per genotype and timepoint). RNA integrity was checked by an Agilent 2100 Bioanalyzer using the RNA 6000 Nano assay (Agilent). All processed samples had a RIN value ≥ 9. The seq library was prepared with a standard TruSeq RNA Sample Prep Kit v2 protocol (Illumina), as described previously (Mariottini et al., 2019). Size and concentration of the RNA-seq libraries was measured by the Agilent 2100 Bioanalyzer using the DNA 1000 assay (Agilent) before loading onto the sequencer. RNA libraries were sequenced on the Illumina HiSeq 2000 System with 100 nucleotide single-end reads, according to the standard manufacturer’s protocol (Illumina). For RNA-seq data analysis, Star 2.5.4b and bowtie2 2.1.0, samtools 0.1.7 and cufflinks 1.3.0 were used as described previously (Stillitano et al., 2017;Hansen et al., 2019;Mariottini et al., 2019). Before read alignment, we identified the sample with the lowest read counts (16,726,861 reads) and down-sampled all other samples to the same number of read counts by randomly removing excessive reads (Supplementary Figure 4A) to prevent distortion of RNA-seq results by read count imbalances, as described previously (Stillitano et al., 2017). The percentage of reads that were successfully aligned to the mouse reference genome was higher than 95% with the exception of one replicate that achieved an alignment success rate of ∼74% (Supplementary Figure 4B). Differentially expressed genes (DEGs) were identified based on a maximum False Discovery Rate (FDR) of 5% and a minimum fold change of log_2_((FPKM_condition1_ + 1)/(FPKM_condition2_ + 1)) > = ±log_2_(1.3) (Supplementary Table 1). Up- and down-regulated genes were submitted to pathway enrichment analysis using Fisher’s Exact Test and the Gene Ontology Biological Processes 2018 library downloaded from enrichR (Chen et al., 2013) (Supplementary Table 2). False discovery rates were calculated based on the Benjamini-Hochberg adjustment. For each list of the up- and down-regulated genes, we ranked the predicted subcellular processes (SCPs) by significance. Significance p-values were transformed into –log10(p-values) and visualized as bar diagrams.

### Statistical Analysis

All statistical evaluations were performed with one-way or two-way mixed model Analysis of Variance (ANOVA), as indicated on the figure legends. Post-ANOVA comparisons were done using either Bonferroni’s multiple comparisons test of significance (for behavior), or Tukey’s multiple comparison test (for biochemistry). All statistical tests were two-tailed and conducted at a 5% significance level using Prism 8 software (Graphpad). Values are expressed as mean ± SEM. *P* values are indicated, when applicable.

## RESULTS

### Generation of an incomplete SCI model using targeted replacement mice expressing human ApoE variants

Studies validating the IH impactor device to produce reproducible contusion SCI in mice have been previously reported (Scheff et al., 2003;Nishi et al., 2007). However, models of SCI using TR ApoE3 and ApoE4 mice have not been characterized. We therefore established a reproducible IH-generated contusion-SCI in these TR mice. Data from the actual impact force and spinal cord displacement using ApoE3 and ApoE4 mice from both genders at similar ages and comparable weights were collected (Supplementary Figure 1A-D). The data was consistent between genders and across ApoE variants. In addition, evaluation of the extent of white matter damage after SCI, measured by fluorescent myelin stain, showed no significant differences between genotypes at the epicenter or up to 300 μm above the epicenter (Supplementary Figure 2).

### Functional recovery is greatly impaired in ApoE4 SCI mice

Functional recovery after a laminectomy (sham controls) or a moderate-severe contusion SCI was evaluated on weekly basis by the BMS open-field test (Basso et al., 2006) starting at 6-days post injury (dpi) and continuing up to 27-dpi (Figure 1A). Sham animals from both genders and ApoE genetic backgrounds showed the maximum possible BMS scores at all timepoints, as expected (Supplementary Figure 3A). ApoE3 and ApoE4 SCI male mice showed similar BMS scores at 6-dpi. However, at 13-dpi, the scores for ApoE3 animals were significantly higher than for ApoE4, reaching a more than one-and-half point difference in functional improvement. These differences were also significant at 20-dpi and 27-dpi with BMS scores again being higher at these timepoints for ApoE3 animals compared to ApoE4 mice (Figure 2A). Similar outcomes were observed in females although changes seemed to be slightly smaller. Differences were not noted at 6-dpi, but starting at 13-dpi, ApoE3 females recovered significantly more than ApoE4 females (Figure 2B). The largest difference was observed also at 13-dpi, with a more than one-and-half BMS point difference between genotypes; scores at 20-dpi and at 27-dpi were also significantly higher for ApoE3 females when compared to ApoE4 females. No significant gender differences were observed between genotypes by BMS test (two-way ANOVA, gender effect, p = 0.38, F(1, 28) = 0.786 for ApoE3, and p =0.83, F(1, 28) = 0.0467 for ApoE4).

**Figure 2.**
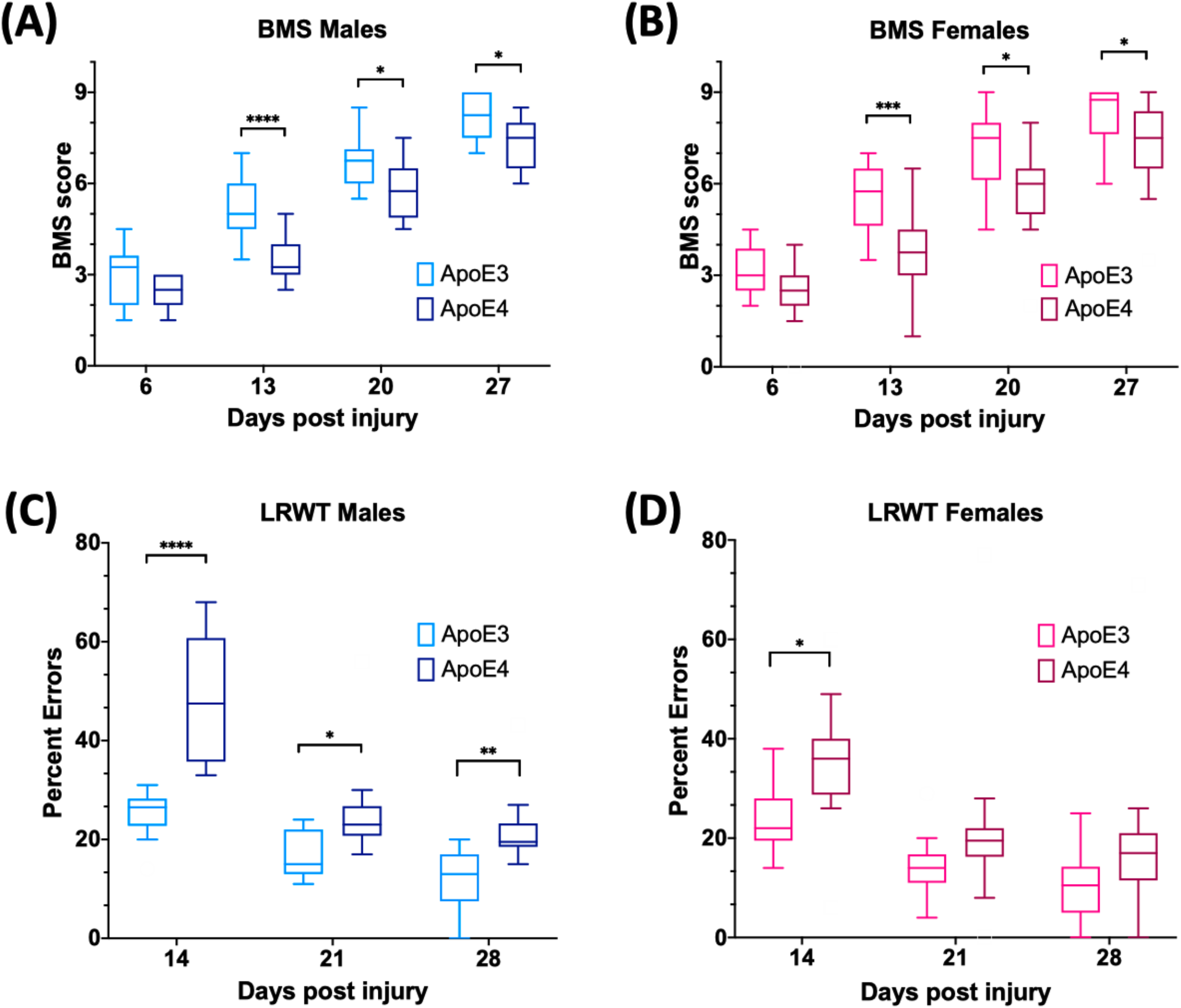
Changes in locomotor function over time for ApoE3 and ApoE4 SCI mice. Weekly BMS evaluation for ApoE3 and ApoE4 (A) male and (B) female mice starting at 6- and up to 27 days post SCI. Weekly LRWT starting at 14 days for ApoE3 and ApoE4 (C) male and (D) female mice. BMS scores range between 0 and 9; LRWT is expressed as percent of foot placement errors. Box-and-whisker diagram represents the median, third quartile (upper box) and first quartile (lower box), and minimum and maximum values (whiskers) of the data. Statistical analysis was performed using a two-way mixed model ANOVA followed by a Bonferroni’s post hoc test. *p<0.05; **p<0.005, ***p<0.001, ****p<0.0001. N = 14 for males and N = 16 for females.

LRWT was performed weekly starting at 14-dpi to complement BMS studies and to evaluate the improvement of fine-motor skills (Cummings et al., 2007). The data demonstrated that while all sham animals behaved like uninjured animals (Supplementary Figure 3B), at 14-, 21- and 28-dpi, ApoE4 SCI male mice had a significantly higher percentage of foot placement errors, which are negative stepping events such as dragging, slipping or missing a rung, compared to animals with the ApoE3 variant (Figure 2C). Females showed similar results with a significant main effect for genotype and time; the only significant pair-wise difference was observed at 14-dpi, albeit a trend toward worse function in ApoE4 mice was also observed at 21- and 28-dpi (Figure 2D). No significant gender differences were observed between genotypes by LRWT (two-way ANOVA, gender effect, p = 0.12, F(1, 30) = 2.61 for ApoE3, and p =0.27, F(1, 26) = 1.26 for ApoE4).

### Differences in neurite sprouting, glial activation and synaptic vesicle markers between ApoE genotypes at the lesion site after SCI

Immunofluorescence (IF) examination of the injured spinal cord at 14-dpi, a timepoint at which changes in functional recovery were maximal between ApoE variants, was performed to assess differences in labeling for proteins associated with neuroregenerative processes such as neurite sprouting, growth cone development and gliosis. A marker for synaptic vesicles was also tested on samples at 28-dpi to investigate changes when locomotor function maximally recovered in both ApoE3 and ApoE4 SCI animals. In addition, these studies were focused on differences at the ventral horn region within transverse sections containing cell bodies of lower motor neurons.

Immunostaining for growth-associated protein (GAP43), known to modulate axonal growth and neuronal plasticity (Koshi et al., 2010), was dramatically increased within ApoE3 sections when compared to ApoE4 at 14-dpi (Figure 3A). Similarly, when compared to ApoE4 mice, an increased expression was observed in ApoE3 mice for 2G13, a growth cone marker associated with the filamentous actin skeleton at 14-dpi (Stettler et al., 1999;Maier et al., 2008) (Figure 3B). When compared to ApoE4 mice, increased staining was also observed in ApoE3 mice for the synaptic vesicle protein synaptophysin at 28-dpi (Figure 3C). Here, IF stained regions were dramatically increased in ApoE3 sections compared to ApoE4. Conversely, detection of glial markers, specifically for reactive astrocytes by staining glial fibrillary acidic protein (GFAP), and macrophage and microglia by staining for ionized calcium binding adaptor molecule 1 (IBA-1), were dramatically increased in sections from ApoE4 mice compared to those from ApoE3 mice at 14-dpi (Figure 3 D,E).

**Figure 3.**
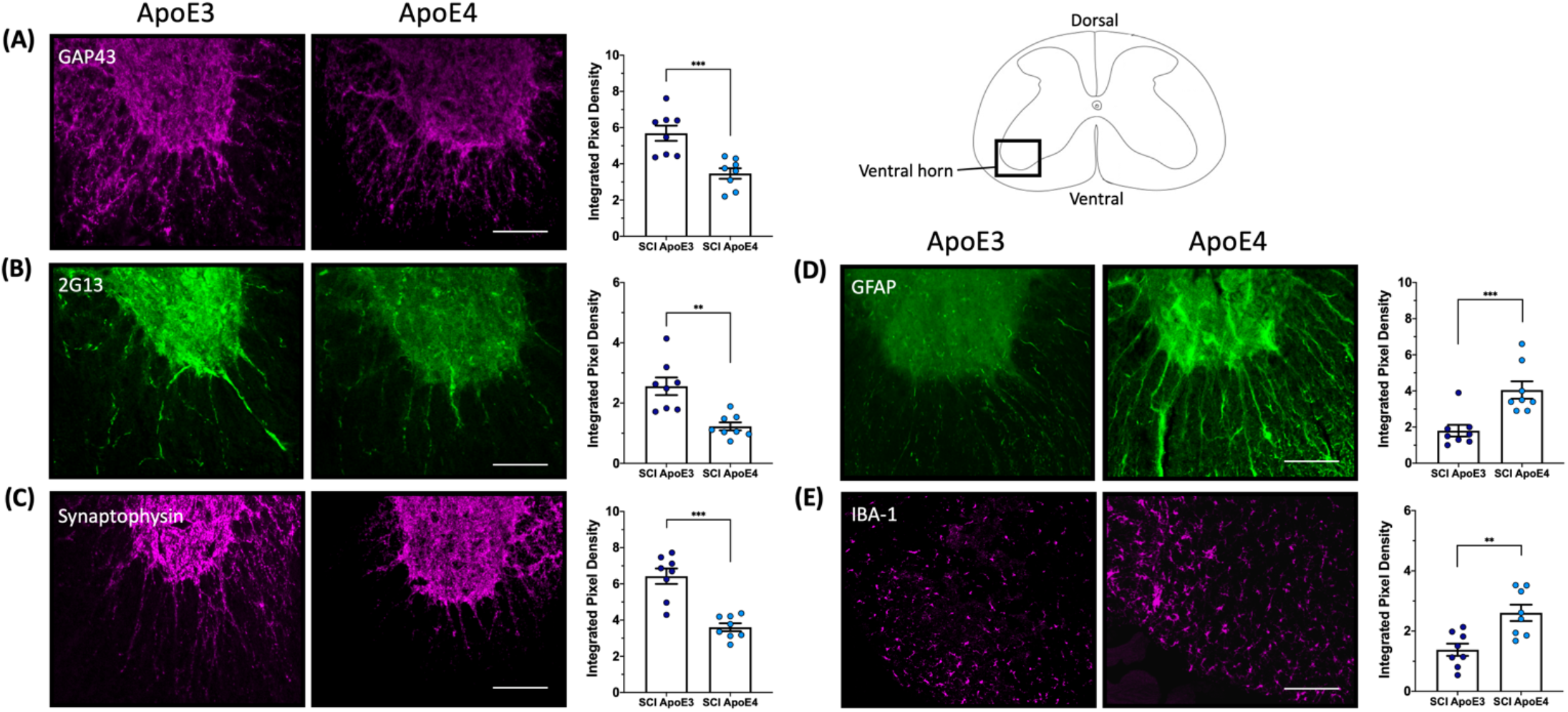
Differences in neurite outgrowth, developing growth cones, presynaptic vesicles, reactive astrocytes and microglia between TR Apo3 and ApoE4 SCI mice. Immunofluorescence staining of transverse 30-micron sections from the lesion site were used to detect: (A) a neurite outgrowth marker (GAP43) and (B) a growth cone development marker (2G13) at 14-dpi; and to label differences in (C) a synaptic vesicle marker (synaptophysin) at 28-dpi. Immunofluorescence staining was also performed to detect (D) a reactive astrocyte marker GFAP and (E) a macrophage and microglia marker IBA-1 at 14-dpi for ApoE3 and ApoE4 SCI male mice. Quantification of immunolabeling by integrated pixel density (right of each panel) was calculated for spinal cord sections and showed significant differences between ApoE genotypes. All images were taken from the ventral horn and are representative examples from N=4 male SCI animals per ApoE genotype. Objective 20X. Scale bar, 100 microns. Bar plots represented as the mean ± SEM (Unpaired t-test followed by Mann-Whitney. **p<0.005, ***p<0.001. N = 8).

### Differences in mRNA levels for genes related to neuroplasticity, astrogliosis and repair between ApoE genotypes after SCI

Dramatic changes in mRNA levels have been reported in the spinal cord after trauma in mice carrying murine ApoE (Liu et al., 2012;Turner et al., 2014;Chen et al., 2015;Mukhamedshina et al., 2017). To test for differences in gene expression between ApoE genotypes, RT-qPCR experiments were run using tissues from ApoE3 and ApoE4 male mice at 14-dpi (Figure 4). mRNA levels of NADPH-dependent diflavin oxidoreductase 1 (Ndor1) were significantly lower in ApoE4 compared to ApoE3 tissues obtained from the rostral half of the lesion site (Figure 4A). Similar changes were observed for neuropilin 1 (Nrp1) (Figure 4B), as well as for the neurite growth marker Gap43 (Figure 4C). In contrast, mRNA levels of matrix metallopeptidase 9 (Mmp9) were higher in ApoE4 compared to ApoE3 tissues from the rostral half of the lesion site (Figure 4D). Several other significant changes were observed within the caudal half of the lesion site. Higher mRNA levels for genes that code for transglutaminase (Tgm2) (Figure 4E) and for astrocytic protein GFAP (Figure 3F) were found in ApoE4 compared to ApoE3 tissues. Conversely, levels of the dopamine transporter gene Slc6a3 (Figure 4G), and synaptosomal-associated protein 25 (SNAP25) (Figure 4H) were found to be significantly lower in ApoE4 compared to ApoE3 tissues obtained from the caudal half of the lesion site.

**Figure 4.**
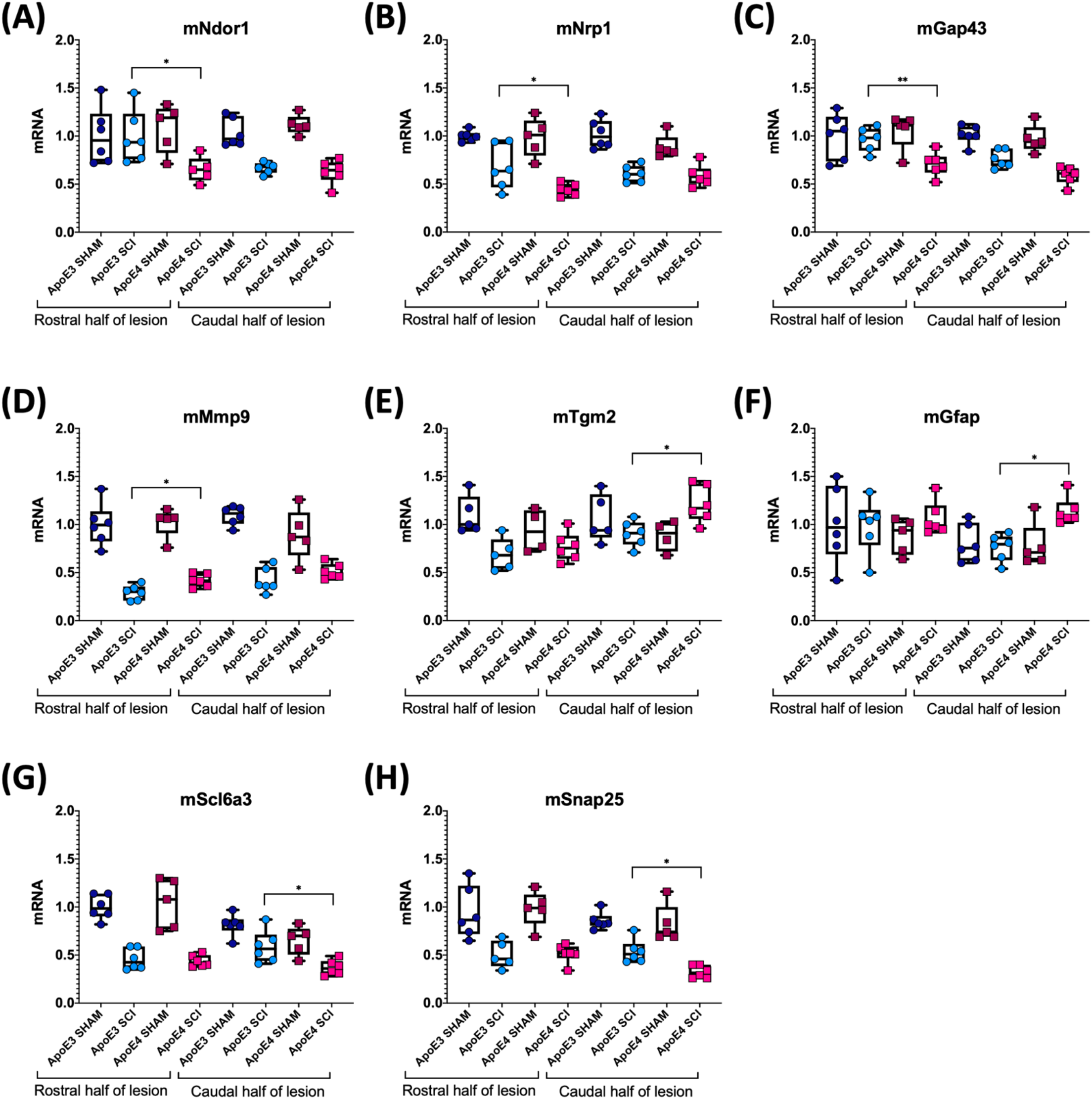
Variations in mRNA levels for genes related to neuroplasticity, synaptic vesicle trafficking, and gliosis between ApoE genetic backgrounds after SCI. RT-qPCR was performed using male ApoE3 and ApoE4 spinal cord tissues at 14-dpi. Levels of (A) Ndor1, (B) Nrp1, (C) Gap43, (D) Snap25, (E) Tgm2, (F) GFAP, (G) Slc6a3, and (H) Mmp9 were detected for laminectomy-only (SHAM), and for SCI animals (SCI) expressing ApoE3 or ApoE4 genotypes. Samples contained either the caudal or rostral half of the lesion site (Figure 1), and data were normalized to each ApoE3 caudal Sham. Box-and-whisker diagram represents the median, third quartile (upper box) and first quartile (lower box), and minimum and maximum values (whiskers) of the data. N=5-6, one-way ANOVA followed by Tukey’s multiple comparisons test *p<0.05; **p<0.005.

### Differences in spinal cord transcriptomic profiles between ApoE3 and ApoE4 SCI mice

A deeper and unbiased understanding regarding the mechanisms responsible for the differences in functional recovery observed between ApoE variants was sought through RNA-seq followed by a bioinformatic analysis. Total RNA was extracted from spinal cord segments at the lesion site (∼4 mm) from male mice at 7-days post SCI (when functional recovery has begun) and 21-days post SCI (when functional differences between genotypes are well-established). Total RNA was extracted from spinal cord segments at the lesion site (∼4 mm) from male mice at 7-days post SCI (when functional recovery has begun) and 21-days post SCI (when functional differences between genotypes are well-established). Differentially expressed genes were identified between both genotypes at each time point. The magnitude of change and number of DEGs that differed between ApoE variants was reported at 7-(Figure 5A) and at 21-dpi (Figure 5B) and showed considerable differences at the latter timepoint, where only 36 DEGs were significantly more highly expressed in ApoE3 mice compared to the 334 that were significantly more highly expressed in ApoE4 mice.

**Figure 5.**
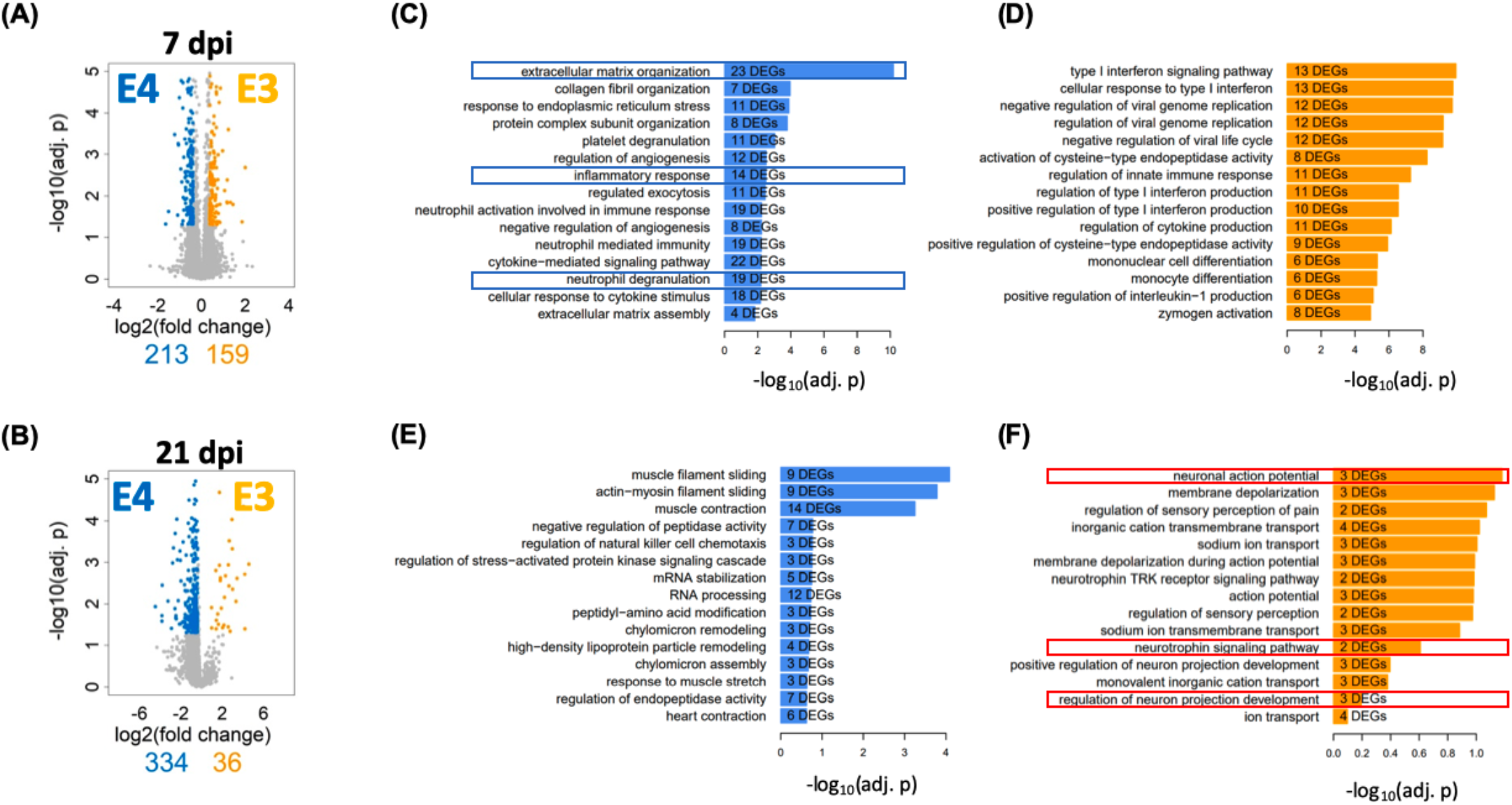
Transcriptome profile at the lesion site for ApoE3 and ApoE4 SCI mice at 7- and 21-dpi. Evaluation of changes in DEGs (FDR 5%, minimum log_2_(fold change) = +/-log_2_(1.3)) between ApoE4 and ApoE3 at (A) 7- and (B) 21-dpi. Colored genes indicate those genes that are significantly upregulated in the indicated condition particular condition, if compared to the other condition. Obtained lists of up-regulated genes were subjected to pathway enrichment analysis using Gene Ontology Biological Process and Fisher’s Exact test, followed by ranking of the predicted pathways by significance. The top 15 ranked pathways are shown for all lists that describe differences between genotypes at both timepoints: (C) ApoE4 (vs ApoE3) at 7-dpi, (D) ApoE3 (vs ApoE4) at 7-dpi, (E) ApoE4 (vs ApoE3) at 21-dpi and (F) ApoE3 (vs ApoE4) at 21-dpi. Numbers of DEGs observed in each particular pathway are shown within the bar for that pathway.

To better understand the biology associated with these sets of DEGs, we identified subcellular processes (SCPs) represented by DEGs that were upregulated in spinal cord tissue from ApoE4 and ApoE3 SCI mice at 7- and 21-dpi and listed these by rank based on adjusted p values (Figure 5C-F). Of note, very different SCPs were found to be upregulated for each genotype and timepoint. SCPs related to Extracellular Matrix Organization, Inflammatory Response, and Neutrophil Degranulation were found for ApoE4 at 7-dpi (Figure 5C). In contrast, for ApoE3 at 7-dpi, the SCPs that were upregulated included functions such as Endopeptidase Activity, Immune Response, and Monocyte Differentiation (Figure 5D). Moreover, at day 21, the SCP for ApoE4 included High-density Lipoprotein Particle Remodeling, and Chylomicron Assembly and Remodeling (Figure 5E), whereas for ApoE3 at 21-dpi the most upregulated SCPs included Neuronal Action Potential, Neutrophil Signaling Pathway and Regulation of Neuron Projection Development (Figure 5F).

## DISCUSSION

The present study explored the role of natural variations of the gene that encodes for human ApoE in determining functional recovery in an animal model after SCI. This novel mouse model provides a clinically relevant starting point for unraveling the mechanisms by which ApoE4 worsens locomotor recovery.

Data from the BMS open-field test indicated that the rate and extent of recovery of locomotor function was lower in ApoE4 animals, regardless of gender, after a moderately severe thoracic SCI compared to ApoE3 mice. In addition, LRWT demonstrated reduced fine motor skills in ApoE4 versus ApoE3 mice. Overall, these differences in behavioral outcomes between TR ApoE3 and ApoE4 mice recapitulate the impaired recovery of function that has been reported in persons with SCI who carry one or two ApoE4 alleles (Jha et al., 2008;Sun et al., 2011).

Given that males account for approximately 80% of patients with SCI (National Spinal Cord Injury Statistical, 2020) and because differences in functional recovery between ApoE genotypes appeared to be largest at 14-dpi, and changes were more evident in males than females, we focused our histological and biochemical studies using tissues at this timepoint. Several kinds of evidence from IF studies of the lesion site support greater neuroplasticity in ApoE3 TR mice. Our histological studies using ApoE3 sections at 14-dpi revealed a greater staining and significant differences for markers of neurite outgrowth and growth cone development by the use of antibodies against GAP43 and 2G13, respectively, as compared to ApoE4. The data suggest that while injured ApoE3 animals demonstrate histologic findings of neuroplasticity, ApoE4 mice either suppresses or confers a loss-of-function for sprouting after SCI. Moreover, we observed a significant increase of the synaptic vesicle marker, synaptophysin, on ApoE3 sections at 28-dpi, a time when substantial locomotor recovery has occurred in this mouse SCI model. Because synaptophysin has been linked to synapse formation and synaptic vesicle trafficking (Tarsa and Goda, 2002;Kwon and Chapman, 2011), these data suggest that not only sprouting but also synapse formation and synaptic vesicle trafficking is higher in injured spinal cords of animals’ carriers of the ApoE3 variant compared to ApoE4.

Conversely, it is well-known that in response to SCI, astrocytes change from a basal state into a reactive state, and that tissue damage leads to the recruitment of inflammatory cells, which at later time points are mainly macrophages and microglia, into the injury site. This process is characterized by morphological changes and functional alterations. Our data demonstrated that appearance of reactive astrocytes, detected by GFAP, was highly and significantly increased at the injury site of ApoE4 samples at 14-dpi when compared to ApoE3. In a similar fashion, staining of macrophages and microglia, when examined by IBA-1, was strongly augmented at 14-dpi in ApoE4 tissues as compared to ApoE3 tissues. The difference in GFAP staining intensity between ApoE3 and ApoE4 tissues is consistent with previous reports that suggest that GFAP has an active role in suppression of neuronal proliferation and neurite extension in the brain, and that astrocytes form a physical barrier to isolate damaged tissue (Brenner, 2014). On the other hand, increased IBA-1 staining in ApoE4 compared to ApoE3 at 14-dpi suggests prolonged and/or greater inflammatory responses and infiltration of immune cells that may be detrimental to the formation of new neuronal connections and sprouting (Wu et al., 2005;Miller et al., 2012).

To complement the histologic findings and study possible differences in mRNA expression of genes related to neuroplasticity and gliosis between ApoE genotypes after SCI, we ran a targeted analysis of mRNA expression using spinal cord segments at the lesion site from sham-operated and SCI male mice at 14-dpi. Consistent with the histological changes reported in ApoE4 samples compared to those from ApoE3, our data confirmed a reduced expression of genes that code for proteins whose functions are related to neuroplasticity through axon growth (Ndor1) (Waller et al., 2017), pruning of corticospinal tract fibers required for motor recovery (Nrp1) (Nakanishi et al., 2019), neurite growth marker Gap43, synaptic vesicle cycling (Snap25) (Antonucci et al., 2016), and dopamine transport via Slc6a3. On the other hand, levels of Mmp9, a matrix metalloprotease involved in extracellular protein degradation (Lu, Takai, Weaver, & Werb, 2011), Tgm2 which is implicated in microglia reparative functions (Giera et al., 2018), and the reactive astrocytic protein GFAP (Brenner, 2014) were significantly higher in ApoE4 samples compared to ApoE3. Taken together, these findings indicate that at 14 days, when functional recovery is rapid and differences in functional outcomes (e.g., BMS scores) between TR ApoE3 and ApoE4 mice are largest, there are clear and substantial reductions in cellular processes that support neuroplasticity in ApoE4 mice associated with increased glial responses.

To further understand the biological basis of the differences in overall function and coordinated stepping between TR ApoE3 and ApoE4 mice, an unbiased approach based on RNA-seq was also employed. Times selected for the RNA-seq analysis were intended to capture events before functional recovery occurred, at 7-dpi, when early neuroplastic changes necessary for such improvements would be anticipated to occur, and at 21-dpi, when functional recovery is well established and neuroplastic responses would be expected to be more focused on strengthening new circuits. This analysis was conducted using ∼4 mm segments of spinal cord tissue containing cells located at the injury epicenter in an effort to capture any differences in biologic responses at the injury site. DEGs were grouped by SCPs to which they contribute. At 7-dpi, the most highly ranked SCPs were involved in inflammatory and immune responses for both ApoE genotypes but, it should be noted, were enriched in genes involved in neutrophil function in ApoE4 mice, whereas genes linked to monocyte and macrophage function were enriched in ApoE3 mice (Figure 5). These changes may reflect a more rapid transition in ApoE3 animals from the initial secondary inflammatory response to injury, which involves infiltration of tissues with neutrophils, to final clearing of debris by macrophages. At 21-dpi, the top-ranked SCP analysis revealed a relative lack of enrichment of SCPs involved in neuronal function in ApoE4 mice, whereas in ApoE3 mice several highly ranked SCPs involved in neuronal action potential, neurotrophin signaling and neuron projection development were identified (Figure 5). These observations suggest more rapid resolution of the inflammatory response in ApoE3 mice and a greater response to support neural circuitry. By contrast, greater or more prolonged neurotrophil infiltration is suggested in ApoE4 mice. Qualitatively, these observations are consistent with the markedly increased evidence of neuroplasticity by histology and targeted analysis of mRNA levels by qPCR observed in ApoE3 mice. The enrichment in subcellular processes involved in neutrophil function observed at 7-dpi in ApoE4 tissues, and suggesting greater tissue inflammation, agrees well with findings from histological studies which indicated a greater degree of activation of microglial cells by Iba1 immunostaining in tissues from ApoE4 mice at 14-dpi, as well as greater immunostaining of activated astrocytes by GFAP (Figure 3).

Taken together, our data suggests that after SCI, ApoE4 mice have worse functional recovery most likely due to reduced sprouting and decreased synaptic transmission compared to ApoE3. Furthermore, ApoE4 injured animals generate a greater astrocytic response and infiltration of microglia and macrophages at the lesion site. Why ApoE4 reduces capacity for sprouting is not known, but several possibilities exist (Toro et al., 2019) (Figure 6). One is reduced efficiency of transport of cholesterol and/or other lipids from astrocytes to neurons, which would lower the availability of cholesterol and lipids for generating new membranes to support axon extension or dendritic projections. Evidence supporting such this mechanism comes from reports showing that ApoE4 is secreted at lower rates than other ApoE variants (Larson et al., 2000;Mishra and Brinton, 2018). In addition, ApoE4 has lower anti-oxidant activity compared to ApoE2 or ApoE3 (Miyata and Smith, 1996) which, coupled with the reduced rate of secretion of ApoE4, would be expected to reduce the net anti-oxidant activity in tissue at the injury site. Recent research on the effects of ApoE variants in the brain further support the concept that ApoE4 impairs sprouting and function. Studies using TR mice demonstrated slower dendritic spine formation and lower spine numbers in ApoE4 mice compared to ApoE2 or ApoE3 mice (Nwabuisi-Heath et al., 2014). Other investigators have reported hyperactivity of specific brain regions by functional magnetic resonance imaging (fMRI) in ApoE4 mice that was attributed to reduced glutamate responsiveness (Nuriel et al., 2017), suggesting that both structure and function of synapses in the spinal cord are influenced by ApoE4. One interpretation of our findings in TR mice with SCI is that the neurotrauma exacerbates these inherent differences in sprouting and synapse formation and synaptic vesicle trafficking, perhaps through prolongation of the inflammatory process.

**Figure 6.**
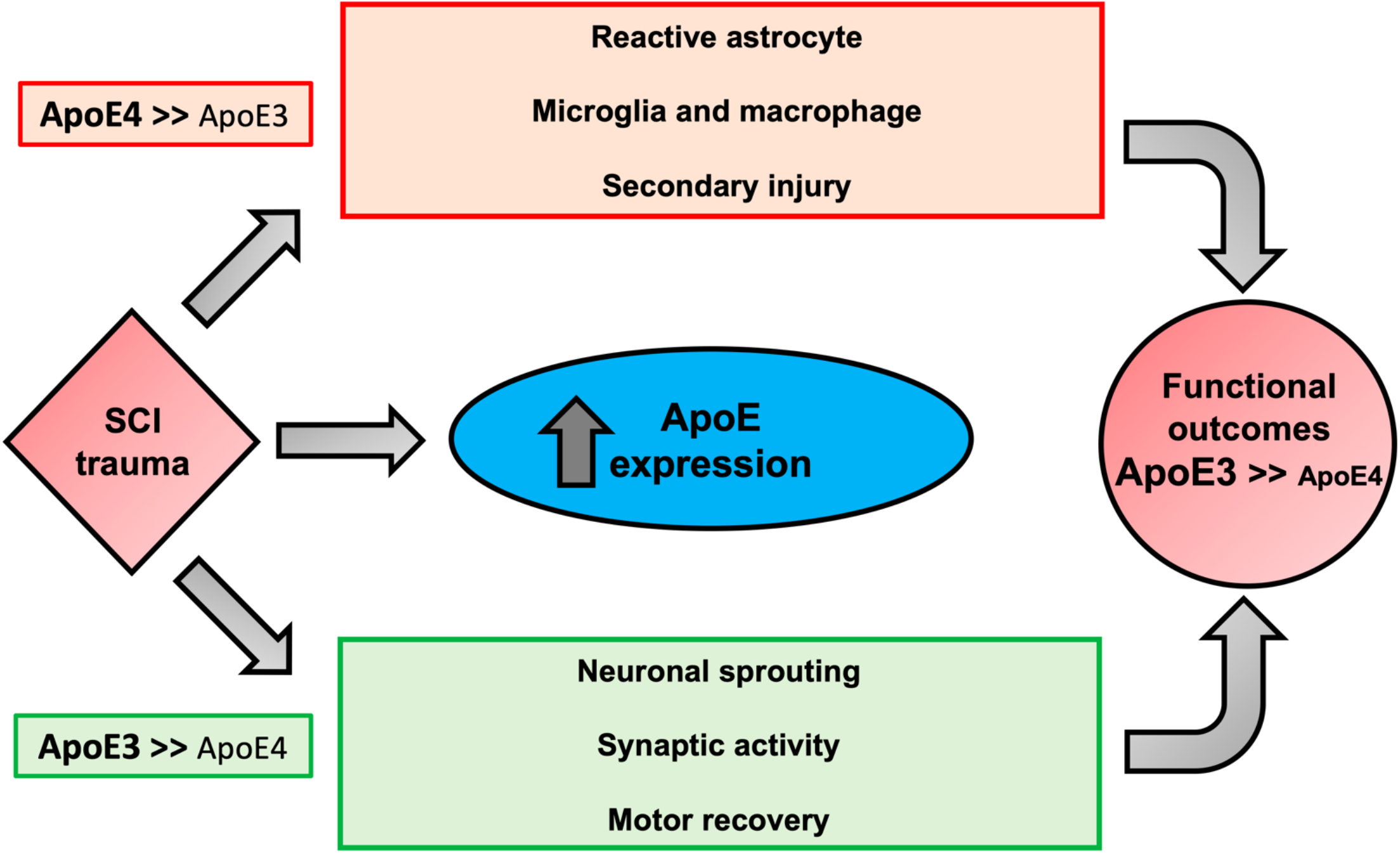
Schematic diagram of functional outcomes after SCI modulated by ApoE genotypes. Proposed mechanisms responsible for the association of the biological activities of ApoE4 to worse neurological outcomes after SCI. ApoE is upregulated in glial cells after trauma. In comparison to ApoE3, ApoE4 is suggested to be less effective in mitigating key factors of SCI pathogenesis, increasing reactive astrocytes and activation of inflammatory cells (i.e. microglia and macrophages), which collectively promote apoptosis and secondary injury due to oxidative stress and production of pro-inflammatory cytokines. ApoE4 also has a lower capacity to stimulate neuronal changes that would be expected to support neuroplasticity and functional recovery such as sprouting, and synaptic activity. In addition, the presence of ApoE4 may diminish transportation from astrocytes to neurons of lipids needed for membrane homeostasis and axon/dendrite growth and may predispose to excess glial scar formation.

## AUTHOR CONTRIBUTIONS

Conceptualization, C.P.C., D.C., and C.A.T.; Methodology, C.A.T, D.D., K.J., D.A., M.S. and J.H.; Software, J.H., M.S., R.S.; Investigation, C.P.C, C.A.T., J.H. and M.S.; Formal Analysis, C.A.T, J.H, K.J, D.D.; Resources, C.P.C, R.I., R.S and D.C.; Writing – Original Draft, C.A.T. AND C.P.C; Writing – Review & Editing, C.A.T., W.Z, M.S., J.H, R.I, D.C, W.B and C.P.C.; Visualization, C.A.T and J.H.; Funding Acquisition, C.P.C., R.I., D.C. and W.B. Supervision, C.P.C., W.B and R.I.

## FUNDING

This work was supported by the New York State, Department of Health, Wadsworth Center, Innovative, Developmental or Exploratory Activities (IDEA) in Spinal Cord (NYS SCIRB Grant# DOH01-PART2-2017-00035 to C.P.C and D.C), the Department of Veterans Affairs Rehabilitation and Development Service Grant B-2020-C, and the James J Peters VA Medical Center. JH, MS and RI were supported by GM54508 and now by GM137056

## CONFLICT OF INTEREST

Authors have no conflicts to disclose, and no competing financial interest exist.

## ACKNOWLEDGEMENTS

We are grateful to Dr. Miguel Gama-Sosa for many helpful discussions about technical aspects of the studies reported herein.

## DATA AVAILABILITY STATEMENT

The data are available upon written, reasonable request. Gene Expression Omnibus (GEO) data accession GSE164688.

## SUPPLEMENTARY MATERIAL

**Supplementary Figure 1.**
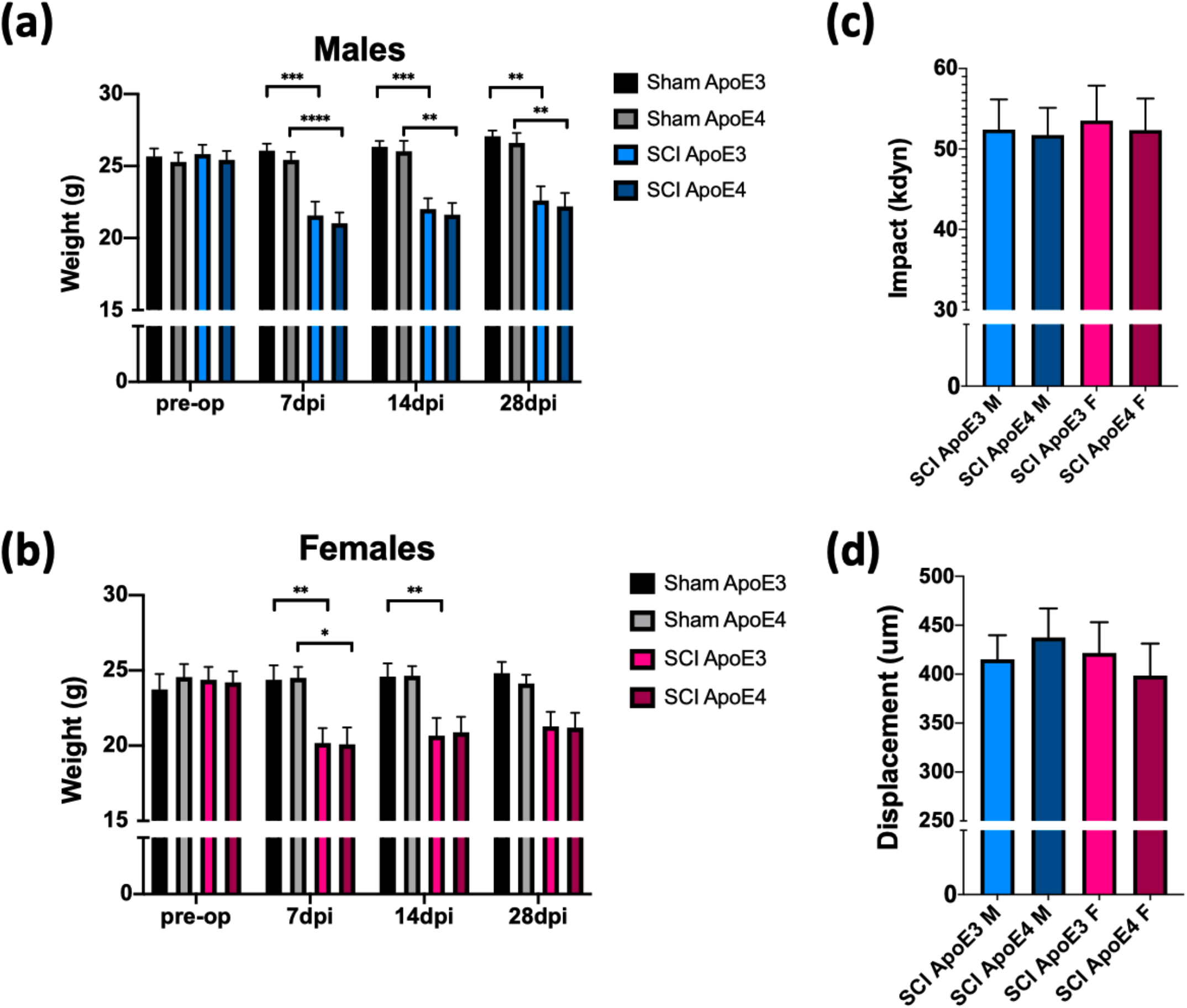
Animal weight, impact force and spinal cord displacement. (a,b) Weight at the time of surgery and weekly thereafter for all experimental ApoE3 and ApoE4 male and female mice. (c) Impact force, and (d) spinal cord displacement recorded by IH device. Bars represent the means plus SEM. Statistical analysis was performed using a two-way mixed model ANOVA followed by a Tukey’s post hoc test. *p<0.05; **p<0.005, ***p<0.001, ****p<0.0001. N=10 for all Shams, N=14 for male-SCI and N= 16 for female-SCI animals.

**Supplementary Figure 2.**
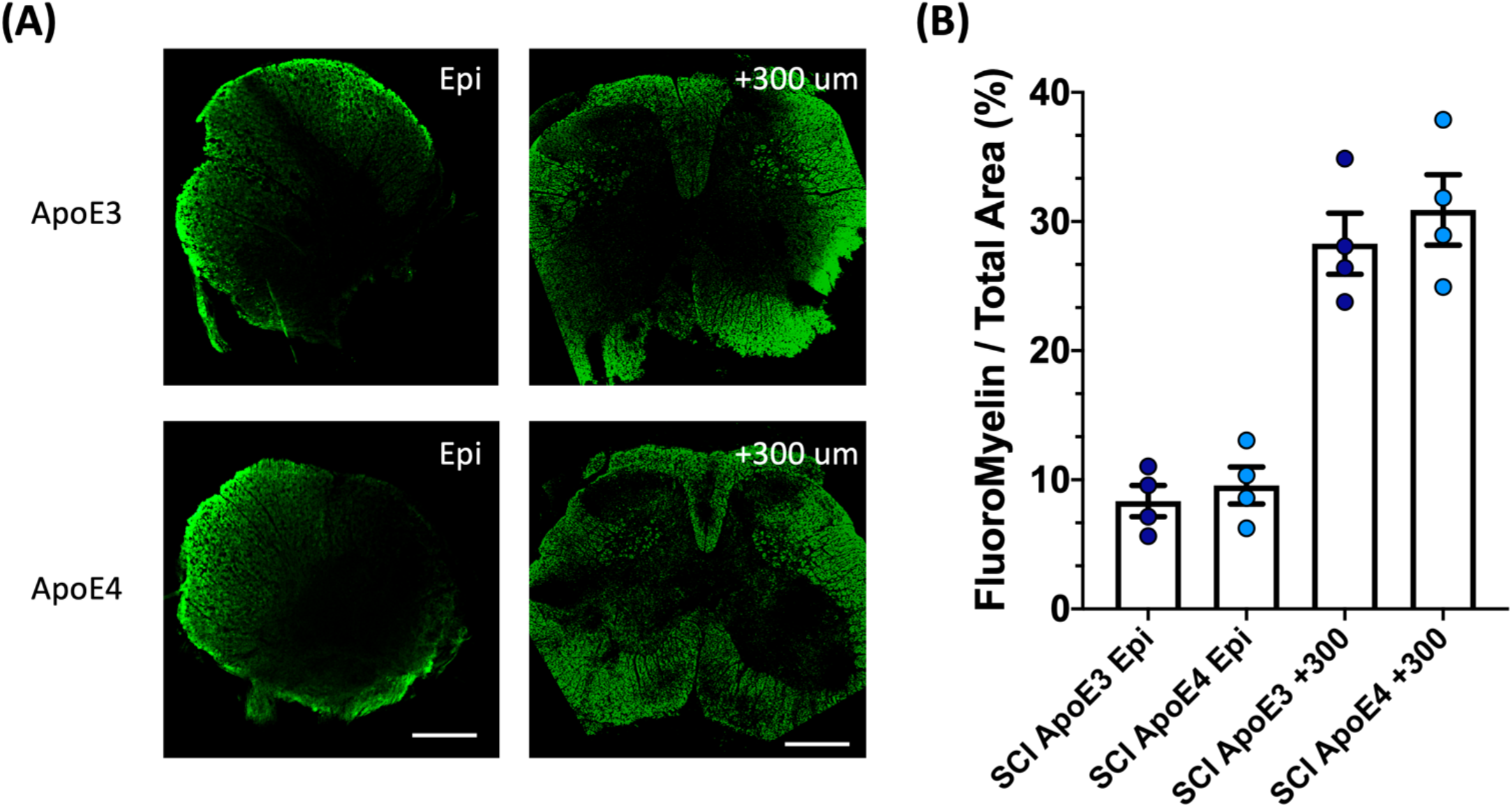
White matter damage in ApoE3 and ApoE4 SCI mice. (A) Extent of white matter damage by detection of fluorescent myelin at the epicenter and 300 microns from the injury site. The scalebar represents 200 microns. (B) Quantification of myelin levels for both genotypes at the epicenter and 300 micros above injury. Bars represent means ± SEM of four mice per ApoE genotype.

**Supplementary Figure 3.**
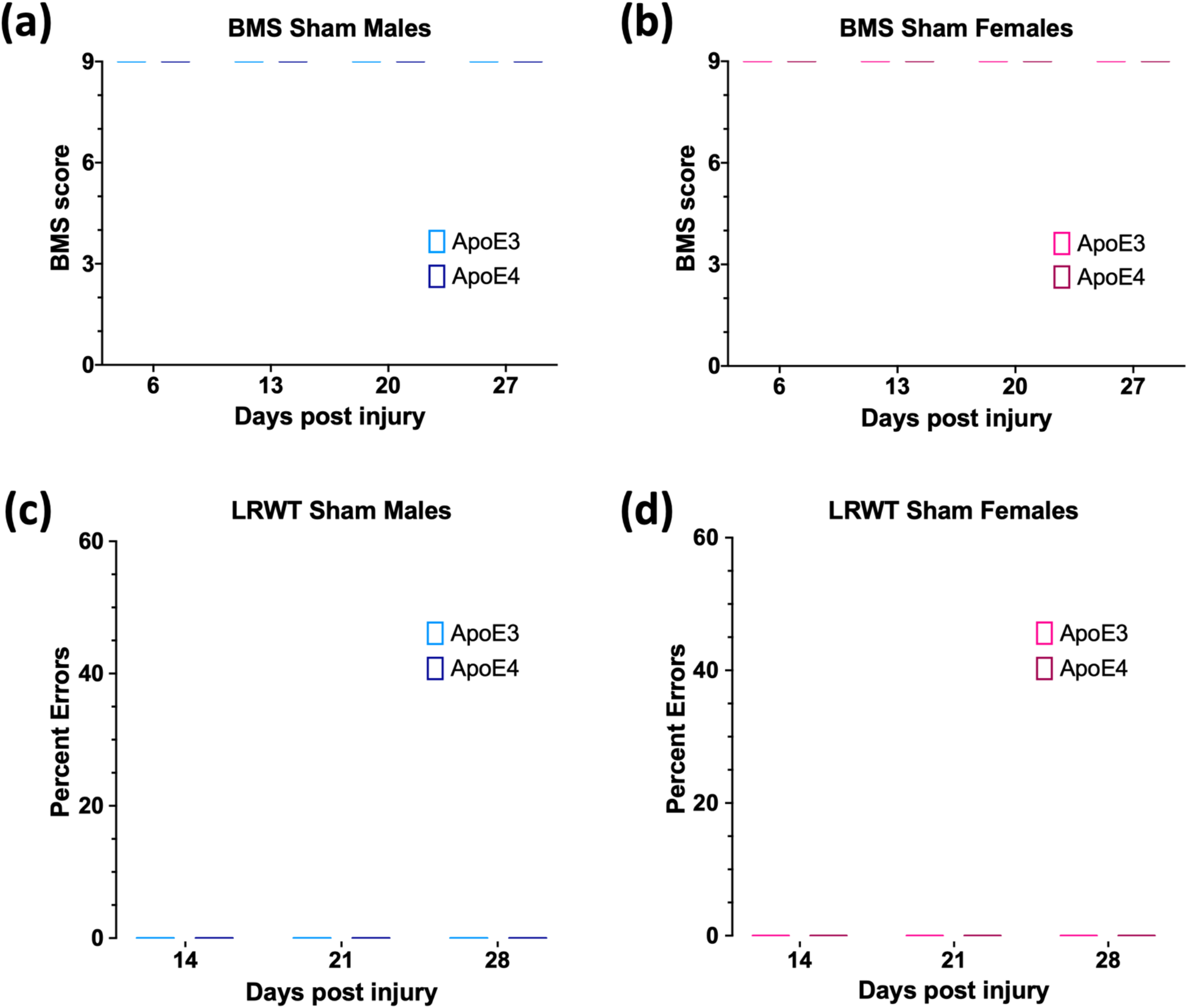
Behavioral test results in ApoE3 and ApoE4 Sham mice. Weekly BMS evaluation for ApoE3 and ApoE4 male (a) and female (b) mice starting at 6- and up to 27 days after laminectomy-only surgeries. Weekly LRWT starting at 14 days for ApoE3 and ApoE4 male (c) and female (d) sham mice. BMS scores range between 0 and 9; LRWT is expressed as percent of foot placement errors. Box-and-whisker diagrams depicting the data are presented. No statistical differences were observed. N=10 for male and female shams.

**Supplementary Figure 4.**
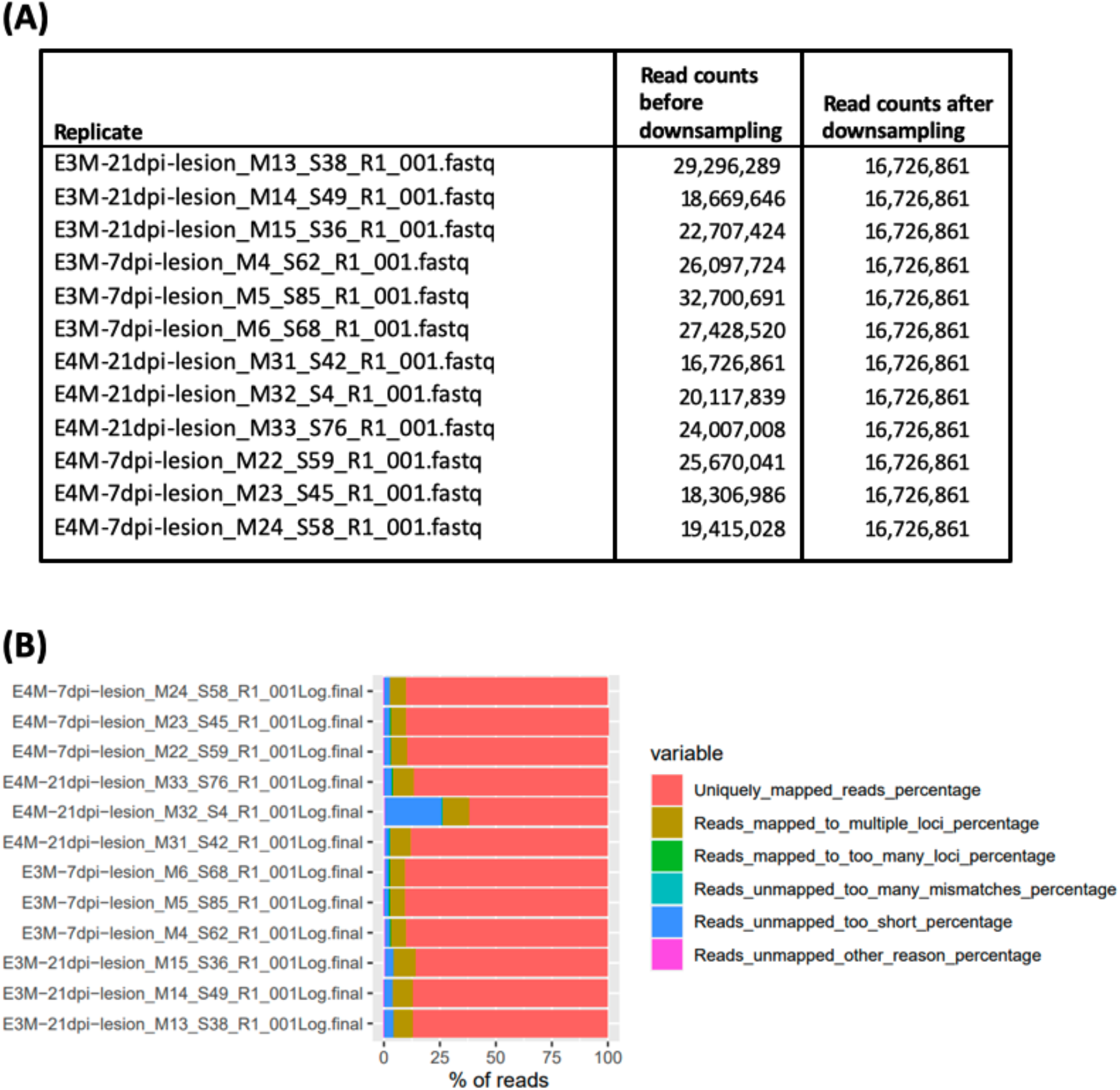
Transcriptomic profiling by RNA sequencing. (A) To avoid distortion of the RNA-seq results due to read imbalances we identified that replicate with the lowest read counts and randomly removed reads from all other samples until they had the same read counts, as described previously (Stillitano et al., 2017). Shown are the read counts of each replicate before and after random read removal. (B) ‘Star’ was used for read alignment and barplots visualize alignment efficiency. For most all except one replicate more than 95% of all reads could be successfully aligned to the mouse reference genome.

<u>Supplementary Table 1.</u> Differentially expressed genes between ApoE4 and ApoE3 mice 7 and 21 days after spinal cord injury (See Excel file attached)

<u>Supplementary Table 2.</u> Predicted top 15 Gene Ontology Biological Processes for each list of differentially expressed genes (<u>See Excel file attached</u>)

## Notes

### Competing Interest Statement

The authors have declared no competing interest.

