## Supplementary figures and images for "The human ApoE4 variant reduces functional recovery and neuronal sprouting after incomplete spinal cord injury in male mice"

### Supplemental Figure 1

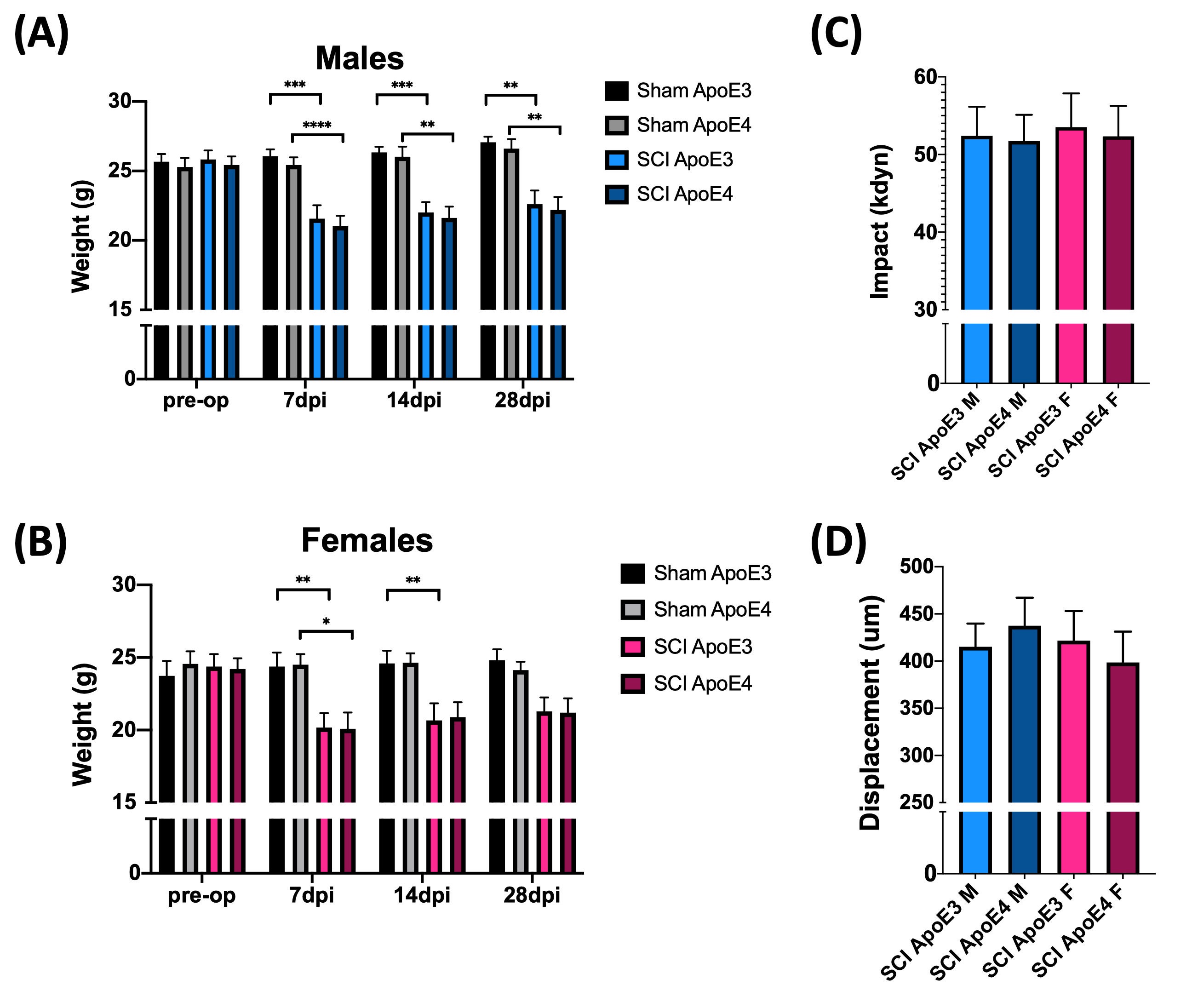

### Supplemental Figure 2

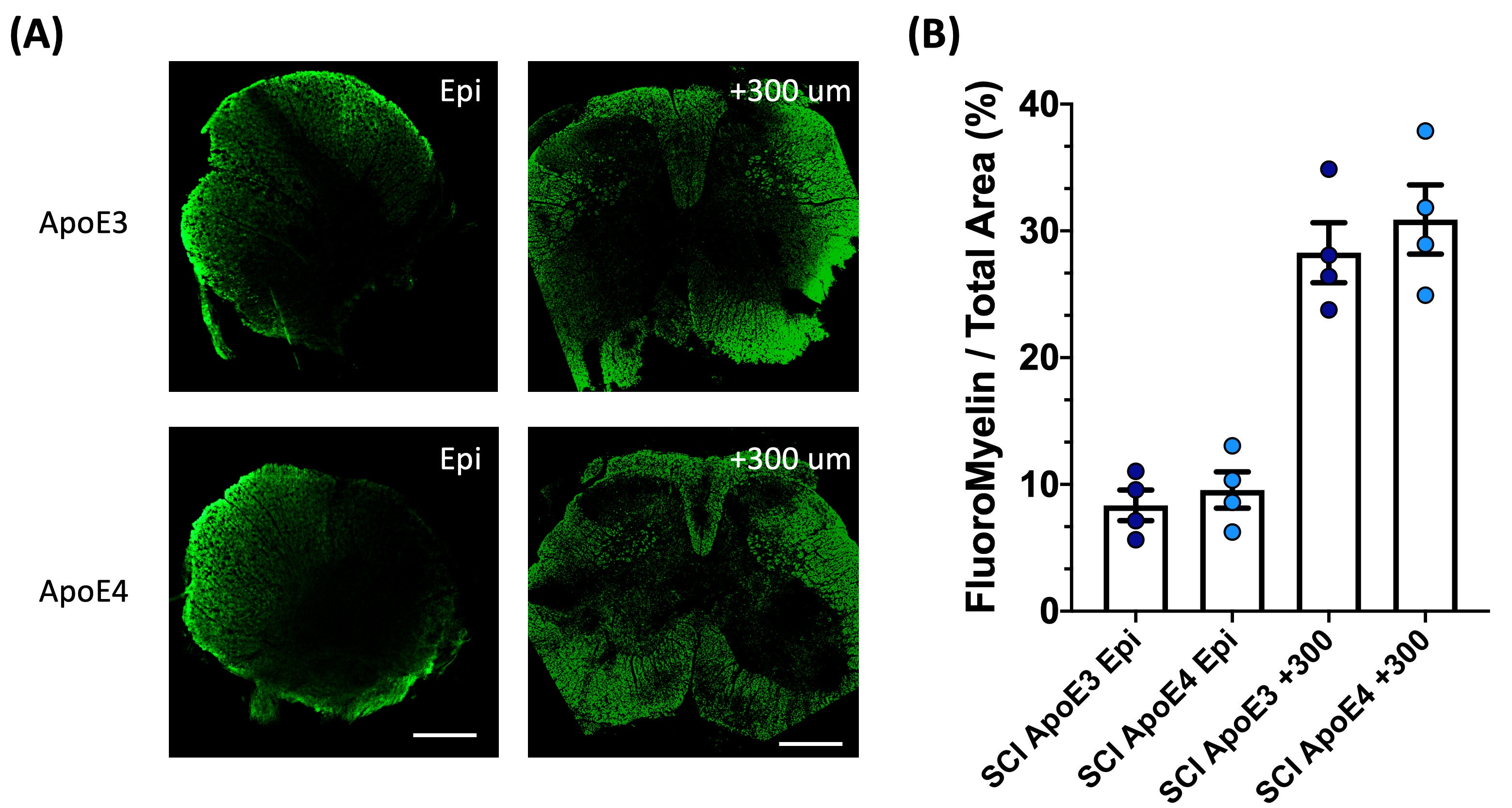

### Supplemental Figure 3

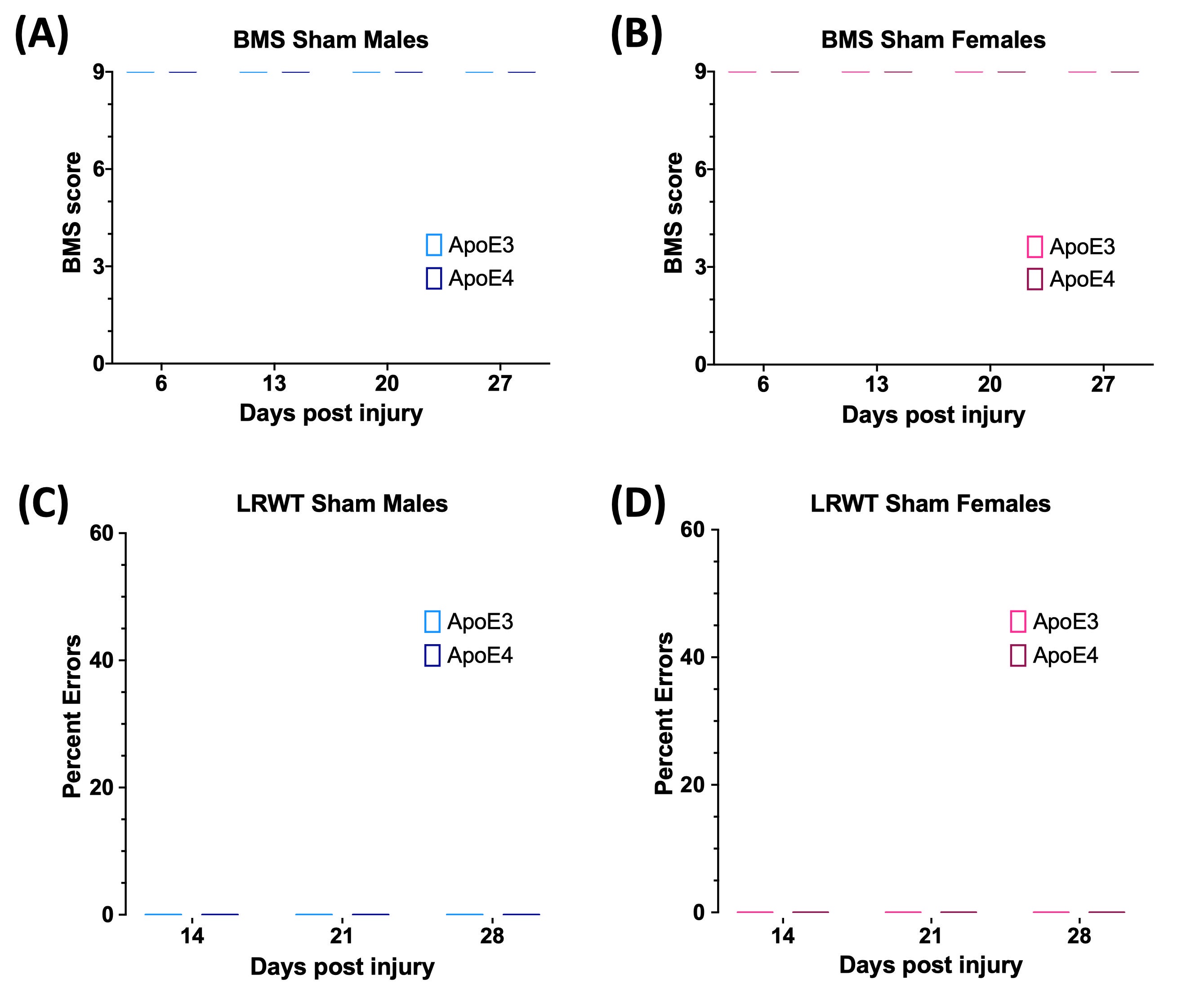

### Supplemental Figure 4

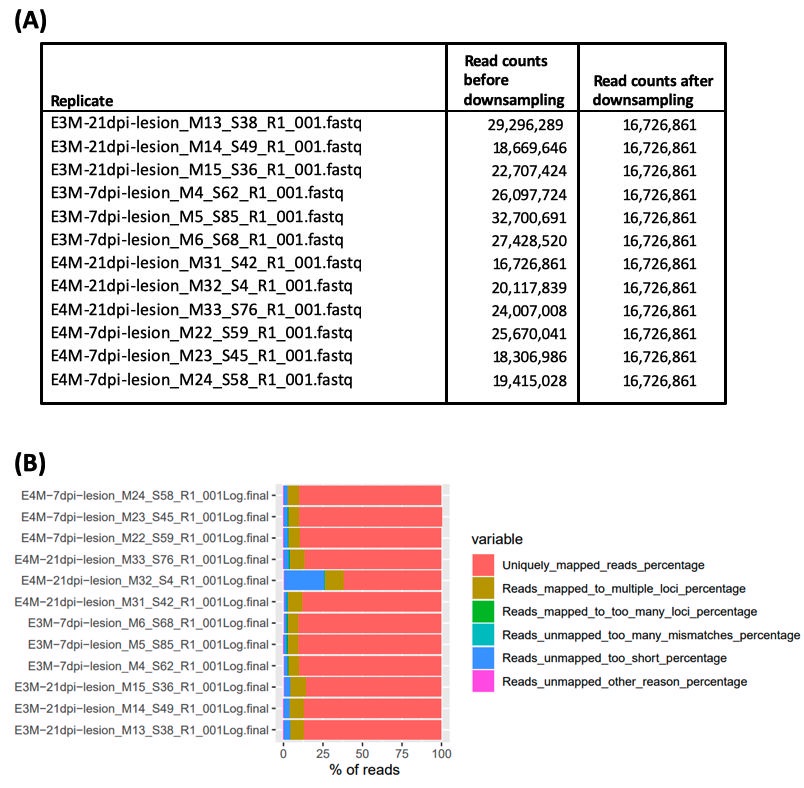
